# Molecular Mechanism of the N501Y Mutation for Enhanced Binding between SARS-CoV-2’s Spike Protein and Human ACE2 Receptor

**DOI:** 10.1101/2021.01.04.425316

**Authors:** Binquan Luan, Haoran Wang, Tien Huynh

## Abstract

Coronavirus disease 2019 (COVID-19) has been an ongoing global pandemic for over a year. Recently, an emergent SARS-CoV-2 variant (B.1.1.7) with an unusually large number of mutations had become highly contagious and wide-spreading in United Kingdom. From genome analysis, the N501Y mutation within the receptor binding domain (RBD) of the SARS-CoV-2’s spike protein might have enhanced the viral protein’s binding with the human angiotensin converting enzyme 2 (hACE2). The latter is the prelude for the virus’ entry into host cells. So far, the molecular mechanism of this enhanced binding is still elusive, which prevents us from assessing its effects on existing therapeutic antibodies. Using all atom molecular dynamics simulations, we demonstrated that Y501 in mutated RBD can be well coordinated by Y41 and K353 in hACE2 through hydrophobic interactions, increasing the overall binding affinity between RBD and hACE2 by about 0.81 kcal/mol. We further explored how the N501Y mutation might affect the binding between a neutralizing antibody (CB6) and RBD. We expect that our work can help researchers design proper measures responding to this urgent virus mutation, such as adding a modified/new neutralizing antibody specifically targeting at this variant in the therapeutic antibody cocktail.

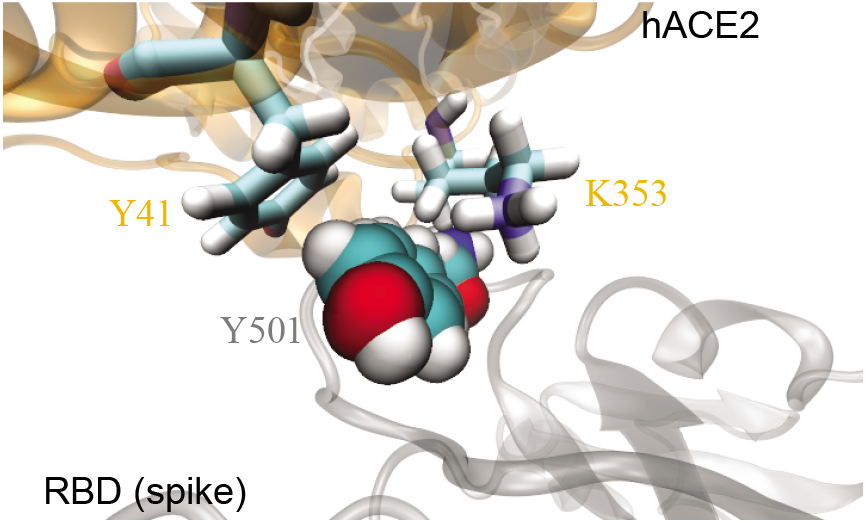

---

The ongoing pandemic of coronavirus disease 2019 (COVID-19) was first detected in China with a cluster of infections caused by a novel coronavirus later identified as the severe acute respiratory syndrome coronavirus 2 (SARS-CoV-2).^1,2^ The disease spreads rapidly across the world with its grave impact reverberating through every corner of the globe, affecting almost every aspect of human life. For example, since the World Health Organization (WHO) declared the COVID-19 outbreak a global pandemic in March, the worldwide COVID-19 confirmed cases has exceeded 80 million with the death toll surpassed 1.7 million in merely nine months. Besides, the global economic recession caused by the COVID-19 pandemic is unusually severe, resulting in a dramatic loss of livelihoods and income on a global scale. Under these unprecedented circumstances, numerous scientists and researchers are racing around the clock to find vaccines and therapeutics to halt the coronavirus pandemic, which has accumulated a wealth of knowledge about SARS-CoV-2.

However, there are growing concerns about the impact of viral genome changes since the D614G alters SARS-CoV-2 both fitness and neutralization susceptibility,^3,4^ and quickly became the dominating variant (>85%) since its first identification in late Jan, 2020. Recently, WHO identified the rapid- and wide-spread of emergent SARS-CoV-2 D614G variant with an additional mutation of N501Y,^5^ which has completely independently emerged in the lineage B.1.1.7 (also known as 20B/501Y.v1) in United Kingdom and in the lineage B.1.351 (also known as 20C/501Y.v2) in South Africa. So far, the molecular mechanism underlying the characteristic N501Y mutation is still enigmatic.

Despite a variety of proteins present in coronaviruses, one such major vaccine and antibody target is the spike glycoprotein (S-protein) of the coronavirus which plays a key role in the receptor recognition and cell membrane fusion process, facilitating viral entry into the host cell.^6–8^ The S-protein is present in the form of trimers on the virion surface forming the distinctive “corona” and is composed of two subunits, S1 and S2. The S1 subunit contains a receptor-binding domain (RBD) which is responsible for virus attachment at the cell surface, while the S2 subunit is responsible for viral cell membrane fusion by forming a six-helical bundle via the two-heptad repeat domain. Previous studies on the pathophysiology of SARS-CoV-2 infection revealed that the human angiotensin converting enzyme 2 (hACE2), an integral membrane protein and a zinc metalloprotease of the ACE family, serves as a high-affinity receptor where RBD of the SARS-CoV-2’s S-protein (referred as sRBD hereafter) binds in order to promote the formation of endosomes to trigger viral fusion activity.^7,9,10^ Importantly, the N501Y mutation occurs at the hACE2’s binding site on sRBD. Previous experiments with adaptation of SARS-CoV-2 in the mouse^11^ and high-throughput screening of all possible mutations in sRBD^12^ have predicted that the N501Y mutation can enhance the binding between sRBD and hACE2.

Here, we are motivated to investigate the molecular mechanism underlying the enhanced hACE2-sRBD binding, induced by the N501Y mutation. Using the all-atom molecular dynamics (MD) simulation as a computational microscope, we directly imaged hACE2-RBD binding at the atomic level, and explored how the N501Y mutation can cause conformational changes for residues residing at the hACE2-RBD interface. Furthermore, we employed the rigorous free energy perturbation (FEP) method to predict the binding affinity difference caused by the N501Y mutation detected in the lineages B.1.1.7 and B.1.351 of SARS-CoV-2.

Figure 1 illustrates the simulation system for modeling the interaction between the hACE2 and sRBD, with the focus on the interfacial region. Detailed simulation protocols are provided in the Methods section. Briefly, atomic coordinates for the complex of hACE2 and sRBD were taken from the crystal structure (PDB: 6VW1). The protein complex was further solvated in a 0.15 NaCl electrolyte. The residue N501 in sRBD locates at the peripheral contact between hACE2 and sRBD (Fig. 1a). During the 185 ns MD simulation, the hACE2-sRBD complex originated from the crystal environment was properly equilibrated in the physiology-like environment. Figure 1b shows the root-mean-square-fluctuations (RMSF) of alpha carbon atoms in the backbone of sRBD. RMSF values for most residues in sRBD are less than or about 1.0Å, indicating that the secondary structure of sRBD was stable. Besides the disordered N-terminal (residues 334-337) and C-terminal (residues 519-527), residues from A475 to N487 (located in a long turn between two short anti-parallel β-sheets near the hACE2-sRDB interface) fluctuated significantly, as manifested through their large RMSF values. However, these residues are not in contact with hACE2 and their fluctuations barely affected the binding stability between hACE2 and sRBD.

**Figure 1:**
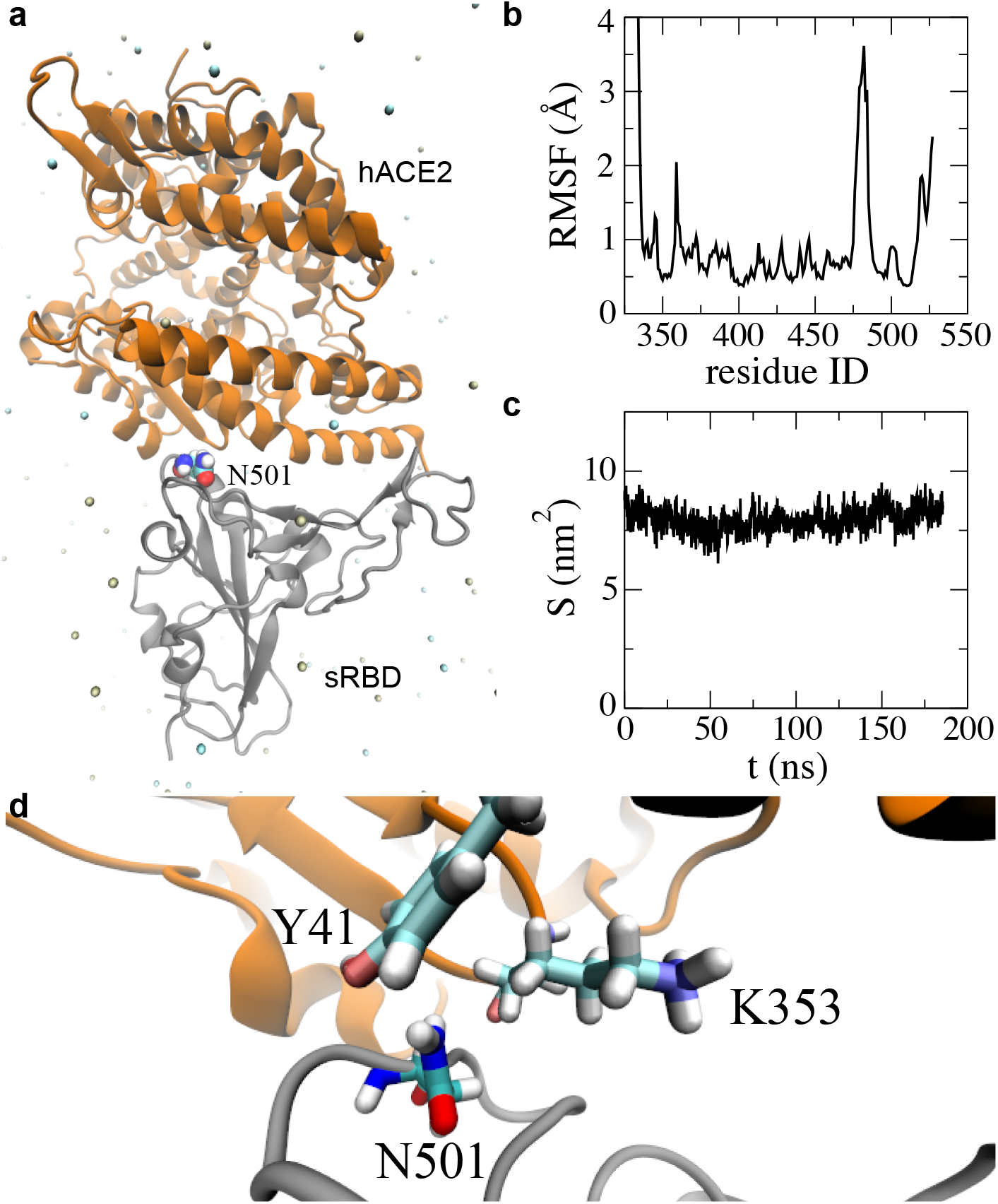
MD simulation of the hACE2-sRBD complex. a) Cartoon illustration of hACE2 (orange) bound with sRBD (gray). The residue N501 on sRBD is in the van-der-Waals sphere representation. Na^+^ and Cl^-^ are shown as tan and cyan balls respectively, and water is not shown for clarity purpose. b) Root-mean-square-fluctuations for residues in sRBD in MD simulation c) Time-dependent interfacial contact areas between hACE2 and sRBD. d) Equilibrated atomic structure for N501 in sRBD coordinating with Y41 and K353 in hACE2.

We then quantified the binding stability by calculating the time-dependent contact area *S* between the hACE2 and sRBD (Fig. 1c). When analyzing the MD trajectory, we first calculated the solvent accessible surface areas (SASA) for both the hACE2 (S_A_) and sRBD (S_B_), and then calculated the SASA for the entire complex (S_AB_). Thus, the contact area S between the hACE2 and sRBD can be estimated as (S_A_ + S_B_ — S_AB_)/2. Figure 1c shows that during the MD simulation, contact areas were nearly constant (~8 nm^2^), indicating a stable binding between hACE2 and sRBD.

By analyzing residues located at the interface, we found that both Y41 and K353 in the hACE2 were within 3.5 Å from N501 in sRBD. Figure 1d illustrates the atomic coordinations among these interfacial residues. During the majority of simulation time, the hydrophilic N501 was in the proximity of the hydrophobic benzene ring of Y41 and the hydrophobic alkane chain in K353. Therefore, it is concluded that these interfacial interactions can be improved with N501 being mutated into a hydrophobic residue. Consistently, through experimental screening of all possible mutations in sRBD, it was found that mutating N501 into V, F, W or Y can enhance the sRBD’s binding with the hACE2.^12^ In the emergent SARS-CoV-2 variant, the presence of the N501Y mutation in sRBD indeed caused the disease to become more contagious, a consequence of the enhanced binding between the hACE2 and sRBD.

To unveil the underlying molecular mechanism of the N501Y mutation, we performed free energy perturbation (FEP) calculations.^13^ As required in FEP calculations, 70-ns-long MD simulations of sRBD alone in a 0.15 M NaCl electrolyte (a free state) was also carried out. After obtaining protein structures for both bound and free states in respective MD simulations, we employed the FEP method to calculate the binding free energy difference for the N501Y mutation on the sRBD, using the thermodynamic cycle shown in Fig. 2. The changes in sRBD’s binding free energies induced by the N501Y mutation can be calculated as ΔΔ*G* = Δ*G*_2_ – Δ*G*_1_ = Δ*GA* — Δ*GB*. In practice, direct calculations of Δ*G*_1_ and Δ*G*_2_ (see Fig. 2) are challenging, which can be circumvented by computing Δ*GA* and Δ*GB* instead. Through the ensemble average,^13^ Δ*GA* and Δ*GB* can be calculated theoretically as 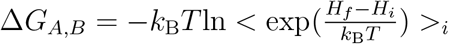, where *k*_B_ is the Boltzmann constant; *T* the temperature; *H_i_* and *H_f_* the Hamiltonians for the initial (*i*) and the final (*f*) stages respectively. As shown in Fig. 2, for the N501Y mutation in bound/free states, the original sRBD with N501 is present in the initial stage and in the final stage the same residue becomes Y501, through the alchemical process in the FEP calculation.

**Figure 2:**
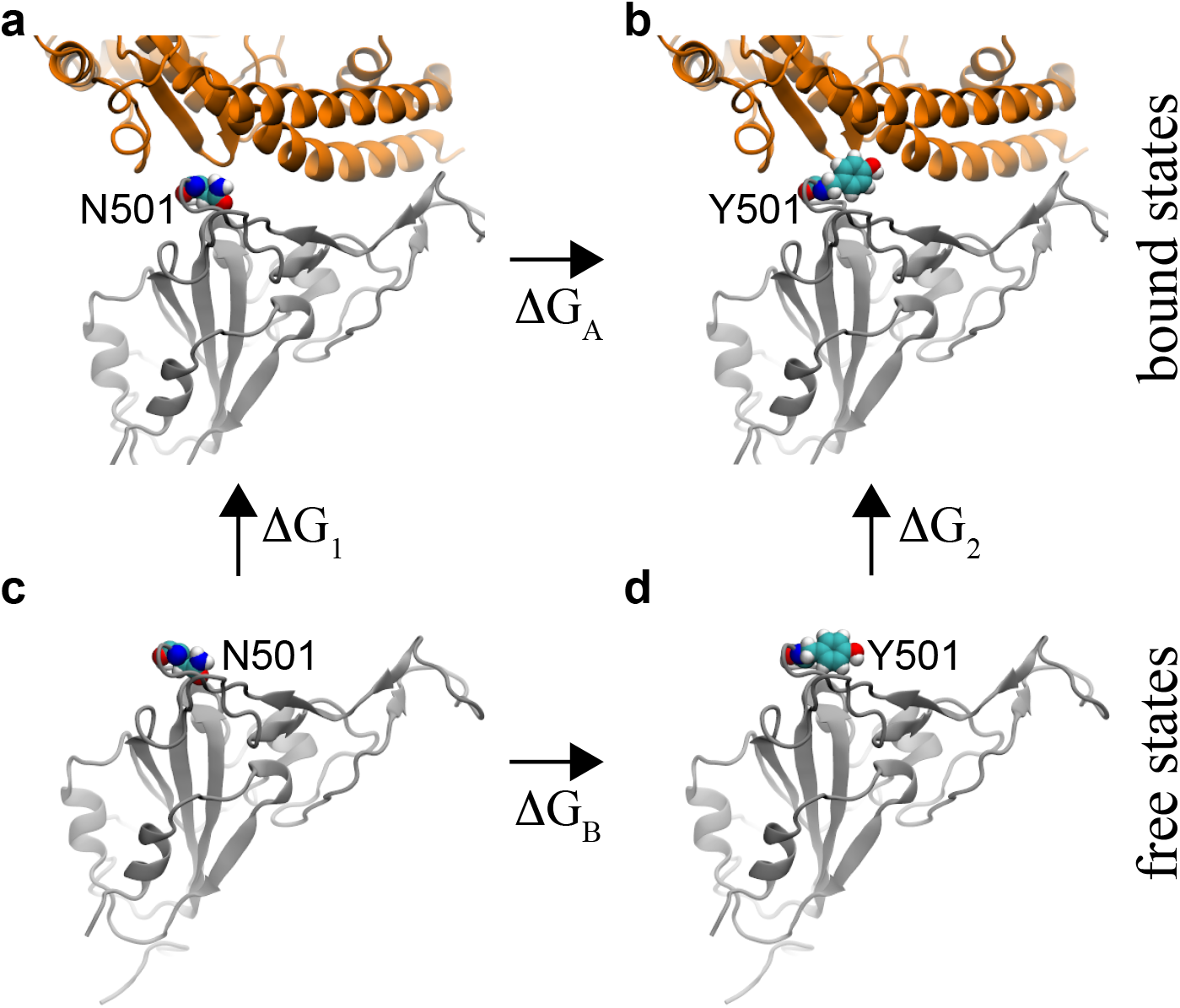
Illustration of a thermodynamic cycle used in the FEP calculations with the mutation N501Y. a) The bound state between the original sRBD and hACE2. b) The bound state between the mutated sRBD (N501Y) and hACE2. c) The free state of the original sRBD (with N501) in water. d) The free state of the mutated sRBD (with Y501) in water. Protein segments (in cartoon representation) are colored the same as those in Fig. 1a.

Table 1 summarizes results from the FEP calculations. In the bound state (Figs. 2a and 2b), the N501Y mutation resulted in an increase in free energies, i.e. Δ*G_A_*=67.19 kcal/mol. In the free state (Figs. 2c and 2d), the same mutation yielded the free energy change Δ*G_B_* of about 68.00 kcal/mol. Note that these energy values for Δ*G_A_* and Δ*G_B_* include the interaction energy change between a residue and its environment as well as the internal energy change within the residue. The latter cancels out when calculating ΔΔ*G*. Overall, the value of ΔΔ*G* is −0.81 kcal/mol, suggesting that the N501Y mutation increases the binding affinity between hACE2 and sRBD (consistent with previous experimental results^11,12^).

**Table 1:** Values of ΔΔ*G* for the N501Y mutation, in sRBD’s binding with hACE2 and CB6 (mAb).

| N501Y | $\Delta G_{A,B}$ (kcal/mol) | $\Delta\Delta G$ (kcal/mol) |
| --- | --- | --- |
| sRBD only | 68.00±0.18 | — |
| sRBD-hACE2 | 67.19±0.65 | -0.81±0.67 |
| sRBD-CB6 | 68.62±0.43 | 0.62±0.47 |

We further explored the molecular mechanism of the N501Y mutation through analyzing the interfacial atomic structures in the simulation trajectory. As mentioned above, in the wild-type S-protein N501 is actually unfavorably coordinated by hydrophobic fragments in Y41 and K353 of hACE2. With the mutation, Fig. 3 demonstrates that the hydrophobic pocket formed by Y41 and K353 (in hACE2) can be fit well by Y501 (in sRBD). Evidently, the edge of Y501’s benzene-ring is in contact with the surface of Y41’s benzene-ring (Fig. 3), forming the perpendicular T-shaped contact which is a well known hydrophobic interaction other than the parallel π – π stacking. Additionally, the surface of Y501’s benzene ring interacts hydrophobically with the alkane chain of the amphipathic K353 (Fig. 3). Therefore, Y501 in sRBD coordinates very well with both Y41 and K353 in hACE2. In Movie S1 (Supporting Information), we show how during the alchemy FEP calculation the exnihilated Y501 (in sRBD) gradually makes a good coordination with Y41 and K353 (in hACE2).

**Figure 3:**
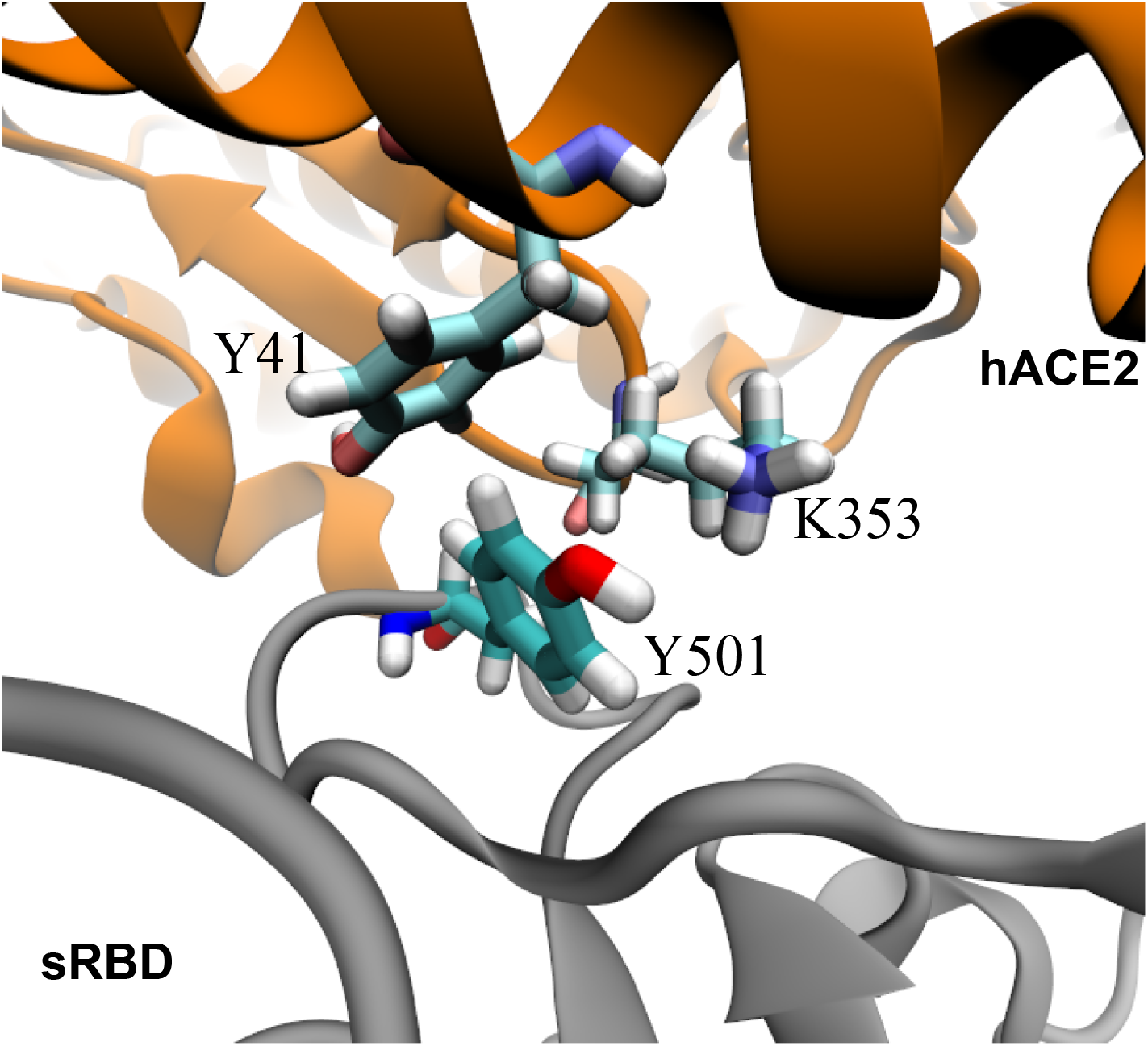
Enhanced interfacial coordinations between Y501 in sRBD and key residues (Y41 and K353) in hACE2.

The above results suggest that the N501Y mutation is favorable in the bound state. On the other hand, in the free state the hydrophilic N501 coordinates surrounding water molecules better than the hydrophobic Y501 does. Thus, the N501Y mutation in the free state is unfavorable. Taking all together, we found that Y501 in mutant sRBD energetically favors the bound state and can enhance the binding affinity between hACE2 and sRBD (i.e. ΔΔ*G*<0).

Currently, neutralizing monoclonal antibodies (mAbs) hold the promise to be both therapeutic and prophylactic for COVID-19. It is still unknown how the N501Y mutation affects the binding between the S-protein of SARS-CoV-2 and neutralizing antibodies. Apparently, this mutation cannot affect the binding of mAbs that target domains other than sRBD, such as the N-terminal domain of S-protein.^14^ Additionally, even targeting sRBD, some mAbs (such as CR3022,^15^ S309^16^ and REGN10987^17^) bind different epitopes on sRBD and thus the N501Y mutation has little effect on those bindings. However, many other mAbs (such as CB6,^18^ P2B-2F6,^19^ B38 ^20^ and REGN10933^17^) bind the epitope site in sRBD that overlaps with the binding site of hACE2. Therefore, the N501Y mutation is likely to affect the binding of these mAbs to sRBD. Below, we focus on how the N501Y mutation can affect the binding between sRBD and the mAb CB6.

The bound state for the complex of sRBD and CB6’s Fab (Fig. 4a) were obtained from our previous 200-ns-long MD simulation. ^21^ The complex highlights that sRBD mainly interacts with the heavy chain (blue in Fig. 4a) in the Fab. Although not as important as the heavy chain, the light chain (orange in Fig. 4a) also contacts the sRBD and the residue N501 is inside this contact (Fig. 4a). From FEP calculations, we found that Δ*G_A_*=68.62 kcal/mol for the N501Y mutation in the bound state. With the value of Δ*G_B_* for the N501Y mutation in the free state (Tab. 1), ΔΔ*G*=0.62 kcal/mol. The positive value of ΔΔ*G* indicates that the N501Y mutation can weaken the binding between sRBD and the CB6’s Fab.

**Figure 4:**
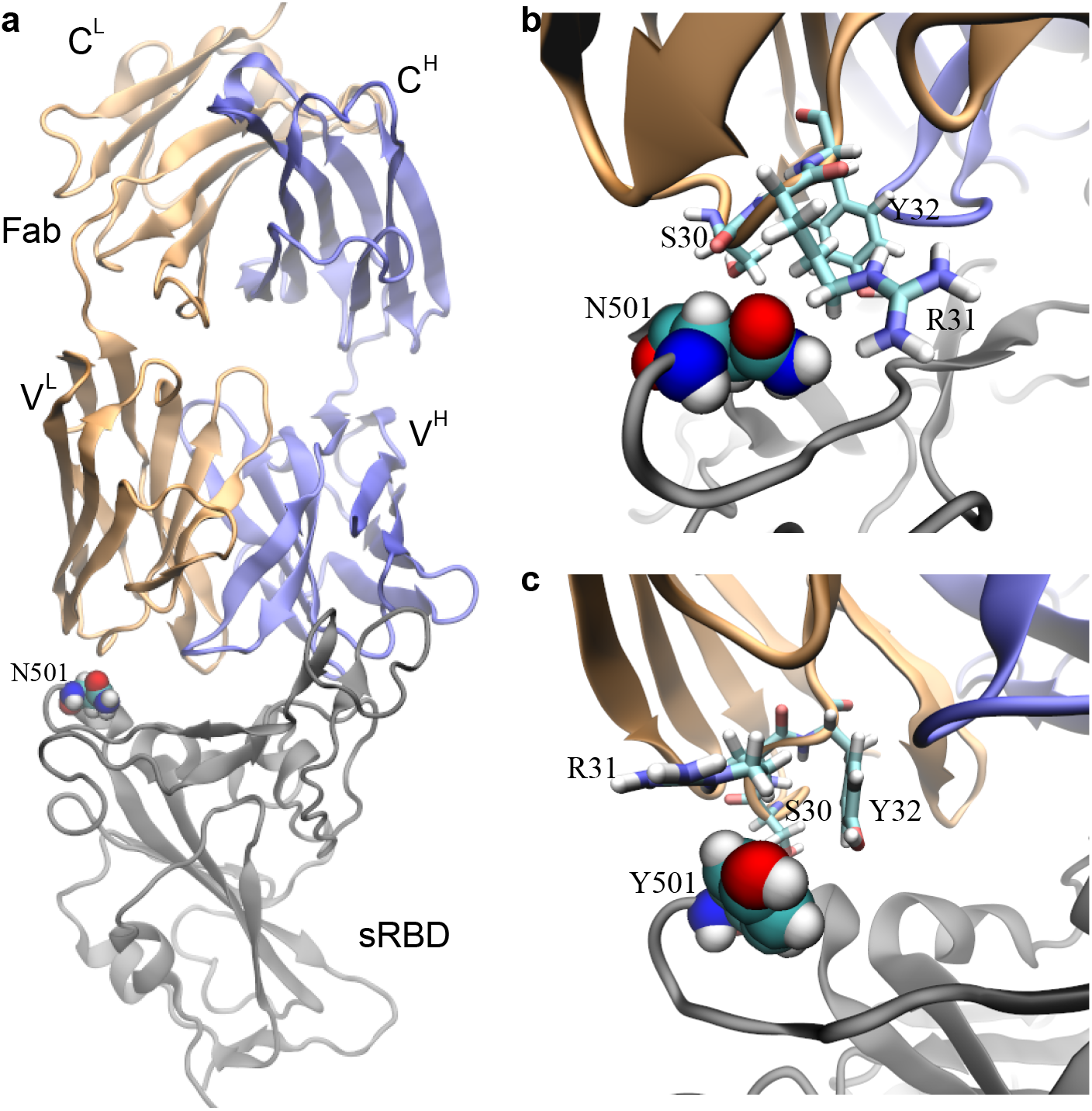
Effects of the N501Y mutation on the binding between the neutralizing mAb CB6 and sRBD. a) The complex of sRBD and the Fab (in CB6) equilibrated in MD simulation. The Fab comprises one heavy chain (fragment) and one light chain, colored in blue and orange respectively; the sRBD is in gray. The heavy (light) chain contains a variable region V^H^ (V^L^) and a constant region C^H^ (C^L^). b) Coordinations between N501 in sRBD and its surrounding residues (S30, R31 and Y32) in Fab, at the beginning of the FEP calculation. c) Coordinations between Y501 in sRBD and surrounding residues (S30, R31 and Y32) in Fab, at the end of the FEP calculation.

Figure 4b illustrates the atomic coordination between N501 in sRBD and the surrounding residues (S30, R31 and Y32) in the Fab. Residues S30 and R31 are closer to N501 than Y32, forming more direct interactions. Despite being hydrophilic, N501, S30 and R31 failed to form stable hydrogen bonds among them. Occasionally, we observed the hydrogen bond between the -NH atoms in the guanidino group in R31 and the oxygen atom in the carboxamide side-chain in N501, indicating a weak attraction. In the final stage of FEP calculations for the N501Y mutation, Fig. 4c illustrates that hydrophobic Y501 repels the charged guanidino group in R31 and is not close to the hydrophobic hydrocarbons in R31. Y32 and Y510 are spatially distant from each other, forming neither the T-shaped nor the π – π interactions. Additionally, S30 is not in close contact with Y501 as well. Overall, Y501 cannot coordinate well with nearby residues from the CB6’s Fab, which accounts for the positive value of ΔΔ*G*.

As a biologics drug, the CB6 can be modified accordingly to accommodate the N501Y mutation in sRBD. According to Fig. 4c, it is recommended to mutate residue 31 in the light chain of the CB6’s Fab from R to F, Y or W, which can increase the hydrophobic interaction across the binding interface.

In summary, our *in silico* studies suggest that the N501Y mutation can enhance sRBD’s binding affinity with hACE2 and potentially cause the virus to evade antibody neutralization. These results are consistent with the fact that N501Y is a naturally occurred and selected mutation.^11^ Prior to July, it had been already discovered that the N501Y mutation could be generated *de novo* from an adaptive murine model using only parental wild-type virus strain (IMEBJ05) first isolated in Beijing.^11^ It only took the parental strain one passage to start exhibiting and accumulating the N501Y variant and by the 3rd passage, the N501Y strain took over 93% of the viral population, suggesting much favorable adaptation to the host than the wild-type strain, presumably due to the enhanced sRBD’s interaction with hACE2. In a more recent study,^12^ it was further validated that N501Y together with N501F, N501W and N501V exhibited enhanced binding affinity between sRBD and hACE2 *in vitro.* The N501Y mutation only requires one mutation (A23063T^5^), whereas the codon requirement for N501 converting to F, W or V requires at least two bases to mutate simultaneously which is much less likely to occur directly from wild-type strain. Therefore, it is much easier for N501Y to occur naturally by evolution and selection.

We discovered that after the N501Y mutation Y501 in sRBD can simultaneously form hydrophobic interactions with Y41 and K353 in hACE2, yielding an enhanced interfacial binding. Namely, the benzene ring’s edge of Y501 forms the T-shaped interaction with the benzene ring’s surface of Y41, and the benzene ring’s surface of Y501 interacts hydrophobically with the alkane chain in K353 (Fig. 3). Based on the molecular mechanism revealed in this study for the sRBD-hACE2 binding (Fig. 3), we hypothesized that mutations of R31 in the light chain of CB6’s Fab into hydrophobic F, Y or W might as well stabilize and improve the interfacial binding between sRBD and the modified CB6 antibody. This modified mAB specifically targeting the SARS-CoV-2 variant might be added into the antibody cocktail to treat all COVID-19 patients.

Overall, our results complement the experimental findings, provide structural insights to assist in evaluating the functional impact of this mutation, and shed light on designing more efficacious antibodies. Apparently, the N501Y mutation in sRBD provides significant edges for virus to proliferate, therefore continued studies on the interaction between sRBD and hACE2 are warranted for development of treatments against COVID-19.

## Methods

### MD simulations

All-atom MD simulations were carried out for both the bound (the complex of hACE2 and sRBD) and free (stand alone sRBD) states using the NAMD2.13 package^22^ running on the IBM Power Cluster. To model the hACE2-sRBD complex (a bound state), we first obtained the previously resolved crystal structure (PDB code: 6VW1)^23^ from the protein data bank and then solvated the complex (with a bound Zn^2+^) in a rectangular water box that measures about 95× 75× 133 Å^3^. 104 Na^+^ and 79 Cl^-^ were added into the system to neutralize the entire simulation system, setting the ion concentration to be 0.15 M (Fig. 1a). The final system containing 96,897 atoms was first minimized for 10 ps and further equilibrated for 1000 ps in the NPT ensemble (*P* ~ 1 bar and *T* ~ 300 K), with atoms in the backbones harmonically constrained (spring constant *k*=1 kcal/mol/Å^2^). During the production run in the NVT ensemble, only atoms in the backbones of hACE2 that are far away from the sRBD (residues 110 to 290, 430 to 510 and 580 to 615) were constrained, preventing the whole complex from rotating out of the water box. We also performed MD simulation for the sRBD alone in the 0.15 M NaCl electrolyte (a free state) using the same protocol.

With equilibrated structures in bound and free states, we carried out free energy perturbation (FEP) calculations.^13^ In the perturbation method, many intermediate stages (denoted by λ) whose Hamiltonian *H*(λ)=λ*H_f_*+(1-λ)*H_i_* are inserted between initial and final states to yield a high accuracy. With the softcore potential enabled, λ in each FEP calculation for Δ*G_A_* or Δ*G_B_* varies from 0 to 1.0 in 20 perturbation windows (lasting 0.3 ns in each window), yielding gradual annihilation and exnihilation processes for N501 and Y501, respectively. We followed our protocol (used in previous mutagenesis studies for optimizing a neutralizing antibody targeting SARS-CoV-2^21^) to obtain the mean and error for Δ*G_A_* and Δ*G_B_*.

We applied the CHARMM36 force field^24^ for proteins, the TIP3P model^25,26^ for water, the standard force field^27^ for ions. The periodic boundary conditions (PBC) were used in all three dimensions. Long-range Coulomb interactions were calculated using particle-mesh Ewald (PME) full electrostatics with the grid size about 1 Å in each dimension. The van der Waals (vdW) energies between atoms were calculated using a smooth (10-12 Å) cutoff. The temperature T was maintained at 300 K by applying the Langevin thermostat,^28^ while the pressure was kept constant at 1 bar using the Nosé-Hoover method.^29^ With the SETTLE algorithm^30^ enabled to keep all bonds rigid, the simulation time-step was set to be 2 fs for bonded and non-bonded (including vdW, angle, improper and dihedral) interactions, and electric interactions were calculated every 4 fs, with the multiple time-step algorithm.^31^

## Acknowledgement

T.H and B.L. gratefully acknowledge the computing resource from the IBM Cognitive Computing Program.

## Supporting Information Available

Movie S1 (N501Y.mpg): shows how during the alchemical process (FEP calculations) Y501 in sRBD gradually shows up and coordinates Y41 and K353 in hACE2. For clarity purpose, the simultaneous gradual annihilation of N501 is not shown.

